# Actin filament bundling by the Arabidopsis VILLIN4 C-terminus involves the basic segment of the IDR linker and is sensitive to temperature and ionic strength

**DOI:** 10.64898/2026.11.02.763837

**Authors:** Stefan K. Clarke, William P. Pedersen, Jake S. Heins, Caleb M. Howerton, Evan J. Curry, Damien L. Mosier, Lynn J. Pillitteri, John M. Antos, Serge L. Smirnov

**Affiliations:** Department of Chemistry, Western Washington University, 516 High Street, Bellingham, Washington 98225-9150, United States; Biochemistry Program, the Ohio State University, Columbus, Ohio, United States; Pfizer, Bothell, Washington, United States; Department of Biology, Western Washington University, 516 High Street, Bellingham, Washington 98225-9160, United States

**Keywords:** Actin filaments, F-actin, villin, actin bundling, Arabidopsis thaliana, NMR, Fluorescent Microscopy, Intrinsically Disordered Region, Bioinformatics

## Abstract

Plant villins are ubiquitous proteins involved in F-actin regulation in all plant organs. Villins consist of folded core and headpiece domains connected by an intrinsically disordered linker. In plant villins these linkers include up to 200 amino acid residues, which is substantially larger than their vertebrate counterparts (∼40 residues). Moreover, the linkers in most types of plant villins exhibit a distinct charge partitioning pattern: a larger basic N-terminus and a smaller acidic C-terminus. The roles of these distinct, sizable and disordered linkers in plant villins are not well understood and are currently under active scrutiny. Here we demonstrate that the basic portion of the linker in *Arabidopsis thaliana* VILLIN4 (AtVLN4) is capable of actin filament bundling. These interactions are also affected by the presence of the adjacent headpiece domain and are altered by temperature and levels of potassium ions, major abiotic regulators that influence plant growth. Based on these findings, we propose that the F-actin bundling capacity of the basic segment in AtVLN4 is responsive to environmental changes and is a uniform property in closely related villins due to high sequence homology across multiple species.

## Introduction

Actin filaments (F-actin) are a dynamic structural component of the eukaryotic cytoskeleton and are essential for maintaining cell shape, in addition to more specialized cellular functions such as cytokinesis, cell-polarity maintenance, organelle movement, locomotion, signaling and vesicle trafficking.(1–6) Villin (VLN) proteins are a well-established family of F-actin regulators in both animals and plants capable of nucleating, capping, severing, and bundling actin filaments.(7–10) In plants, these proteins play critical roles in root hair development, pollen tube growth, and are responsive to environmental stress.(9, 11–16)

Plant villins have a domain structure similar to their vertebrate counterparts, consisting of an N-terminal core of six gelsolin-homology domains connected via an intrinsically disordered region (IDR linker) to a C-terminal headpiece (HP) domain (**Fig. 1A**).(17) Both of these domains contain dedicated actin binding sites allowing villin proteins to bind multiple filaments and promote dynamic actin bundling. (18, 19) Plant and animal villins share significant primary structure identity (30%-50%) in the core and headpiece domains. They also share substantial similarity in tertiary structure (RMSD within 2.6 Å) and comparable capacity for specific binding of F-actin (K_D_ in 3-7 μM range).(20) However, plant villins contain much larger and feature-rich linkers that have distinct amino acid composition patterns (**Fig. S1A**). Recent data and our preliminary analyses reveal that most plant villin linkers are 3-5x larger (130-200 residues) and have more amino acid charge-partitioning or segregation when compared to their vertebrate analogs (**Fig. 1**, **Fig. S1A**). (20, 21) The functional relevance of these difference remains an area of active investigation where recent work has indicated that a subsection of the AtVLN5 linker fused with the core has actin bundling capacity.(22) We propose that the unique features of plant villin IDRs and their sequence conservation across diverse plant species (**Figs. S1B, S1C**) is highly suggestive of unique biochemical and biophysical properties that are not found in vertebrate villin analogs.(17, 21)

**Figure 1.**
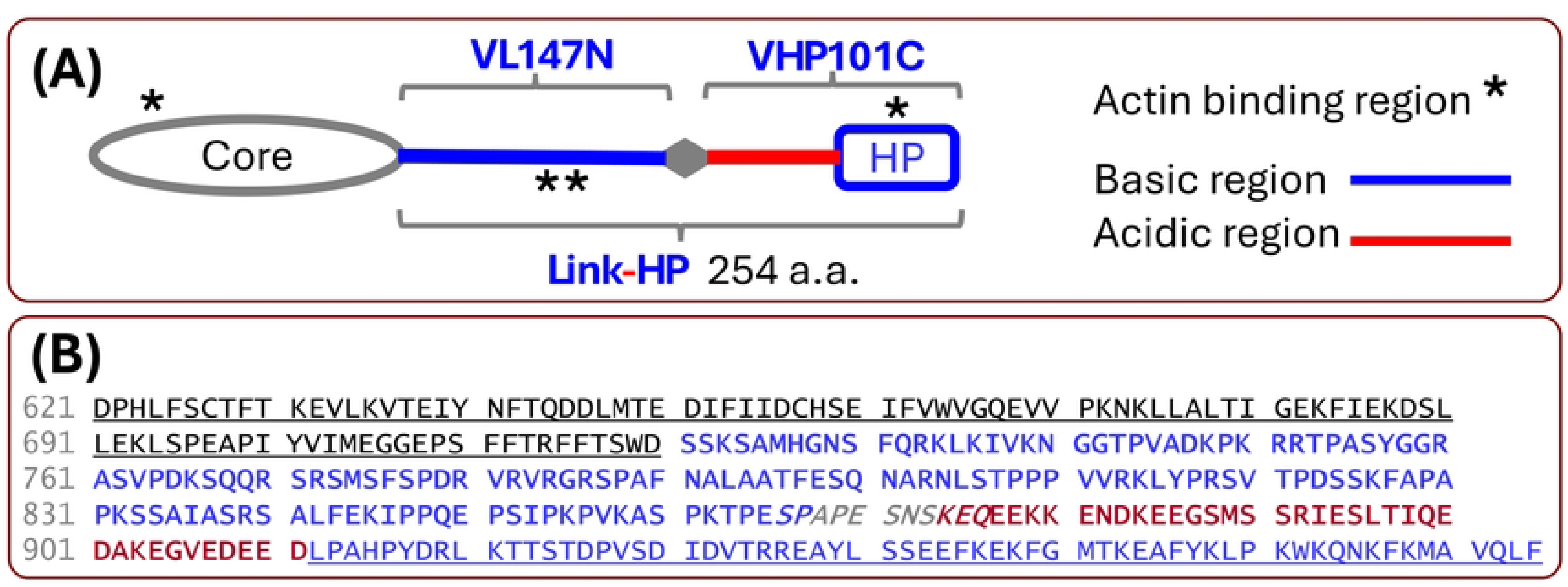
Structural schematic of AtVLN4 and constructs used in this study: VL147N, VHP101C, Link-HP. **(A)** The core and headpiece (HP) domains of *A. thaliana* VILLIN4 isoform 2 (AtVLN4) are connected by a disordered linker.(20, 21) Fragments of AtVLN4 used in this work include are detailed in **Fig. S2**: VL147N (basic portion of the linker, pI 11.7), VHP101C (acidic portion of the linker fused to the headpiece domain, pI 4.1), and Link-HP (full-length IDR linker fused to the headpiece domain). The location of the proposed novel F-actin bundling capability of the AtVLN4 linker is indicated with “**”. The grey rhombus separating VL147N and VHP101C is a predicted regulatory PEST motif (positions 866-876). **(B)** Amino acid sequence of the C-terminus of AtVLN4. This includes core domain 6 (residues 621-720, predicted by BlastP(23) by sequence similarity to the analogous domain in *G. gallus*, PDB 2LLF(24), black underlined), and the C-terminal headpiece domain (residues 912-974, PDB 5VNT(20), blue underlined) connected via the IDR linker (residues 721-911, bold font). Similar to panel **A**, the basic, acidic, and PEST motif portions of the IDR linker are colored blue, red, and gray, respectively. In this work, the bulk of the PEST motif (SPAPESNS) between the basic and acidic segments has been mutated to AAAGGGAG to prevent premature sample degradation, which is the strategy which was successfully implemented for the sizable IDR in the related vertebrate protein dematin.(25)

In considering possible roles for the extended IDR in plant villins, it is important to note that plants generate their organs post-embryonically and are completely sessile organisms that must adapt to various long-and short-term abiotic stresses during growth and development. To this end, plants have a high level of phenotypic plasticity and have evolved multiple strategies to adjust their development in response to both intrinsic and extrinsic cues.(26–28) For example, the actin cytoskeleton is responsive to several stress conditions including extreme temperatures and nutrient-deficiency.(29–31) Arabidopsis mutants defective in the actin binding protein Arabidopsis actin depolymerizing factor 5 (ADF5) have reduced resistance to low-temperature stress, highlighting the importance of the actin cytoskeleton in low-temperature resilience in plants.(30, 32). Under potassium-deficient conditions, plants also modify F-actin dynamics to increase root hair length to improve K^+^ absorption. As noted above, the villin family of proteins have emerged as key regulators of actin remodeling through assembling, depolymerizing, and bundling of F-actin, which is an established mechanism plants use to regulate cellular processes. (26, 33, 34) Recently, Li et. al determined that K^+^-dependent actin modification was associated with an increase in villin bundling function leading to an extension of the elongation phase.(29) Thus, there is accumulating evidence of the role of villin proteins in modulation of the actin cytoskeleton, perhaps by enabling responsiveness to environmental cues such as temperature and nutrient fluctuations, which would be critical for plant survival in a constantly fluctuating environment.(35–38)

In this paper, we describe progress toward elucidating the role of IDRs in plant villins. First, we present a novel software tool (idCHAR) for assessing the extent of charge segregation and the border between the basic and acidic segments within polypeptide sequences and use it to analyze the five villin variants from *Arabidopsis thaliana*. Following this analysis, we experimentally characterized the actin bundling capabilities of select fragments of the IDR linker region in AtVLN4 (VL147N, VHP101C and Link-HP, **Fig. 1**). These studies revealed that the ∼150-residue basic part of the AtVLN4 linker has the novel capacity to bundle actin filaments, which was confirmed by low-speed co-sedimentation with F-actin and fluorescence microscopy. In addition, we demonstrated that actin bundling by this basic linker fragment was affected by the presence of the adjacent headpiece domain and is also sensitive to abiotic environmental changes (K^+^ concentration and temperature). Finally, we utilized solution NMR probing of AtVLN4 fragments at different temperatures and ionic strength as an initial investigation of backbone structure and dynamics. Overall, this work supports a key role for the AtVLN4 IDR linker in modulating actin dynamics and suggests similar properties in other plant villin isoforms.

## Materials and Methods

### Bioinformatics analysis of charge segregation protein sequences

An established bioinformatics tool CIDER(39, 40) used in this work (**Fig. S3A**) was augmented by new software idCHAR which reports specific positions within a polypeptide that separate the basic and acidic segments. Specifically, for every residue position within the target sequence, the code calculates: a) the difference between the number of basic (R, K) and acidic (D, E) residues N-terminal to the sampled position, b) the same value for residues C-terminal to the sampled position and c) the idCHAR score: the difference of the quantities obtained in a) and b). If the values in a) and b) have opposite signs, the position segregates clusters of opposite charges and is analyzed further as follows: a positive idCHAR score indicates a basic N-terminus and acidic C-terminus (+/- scenario); a negative value indicates the opposite (-/+ scenario). The boundary between the two charged segments is identified as the sequence position(s) reporting the greatest absolute value of the idCHAR score. An example of idCHAR score calculation is shown in **Figs. S3B, S3C**. The software idCHAR with its source code can be obtained from https://github.com/ssmirnov56/idCHAR.

### Construction of the VL147N, VHP101C and Link-HP Expression Vectors

The VL147N fragment was designed to include the N-terminal 147 residues of the linker from *AtVLN4* (REFSEQ accession NM_119162.5, UniProt accession O65570). VHP101C encompasses 101 C- terminal amino acid residues of the protein: the acidic part of the linker fused with the headpiece. Finally, the Link-HP fragment encompasses 254 C-terminal amino acid residues of the same protein comprising the linker fused with headpiece. The predicted PEST motif in the linker region was mutated (SPAPESNS to AAAGGGAG) to limit premature protein degradation.(25) All three samples were designed as fusion proteins equipped with an N-terminal affinity tag system: dual Strep and a 10xHis (or 8xHis for VHP101C) tags followed by spacer residues and a TEV protease site. The constructs were cloned into the pET-24a vector (Novagen) using commercial services (Genscript) with primers containing an NdeI site at the 5′ end and a HindIII site at the 3′ end of the TAG stop codon. The amino acid sequences of the three fragments, VL147N, VHP101C, Link-HP, are shown in **Fig. S2A**.

### Protein Expression for VL147N, VHP101C and Link-HP

The standard expression procedure was applied with the BL21(DE3) bacterial expression system transformed with the pET24a vectors and induced with IPTG for protein expression. The overall procedure used was identical to those employed previously. (20, 21, 24, 41) For expression of the fragments labeled with ^15^N isotopes, the cells were transferred to the minimal M9 media augmented with ^15^NH_4_Cl (source of ^15^N) and glucose (source of ^12^C), equilibrated, and induced with IPTG as described before. (20, 21, 24, 41)

### Protein Purification for VL147N, VHP101C and Link-HP

Cells were harvested by centrifugation (10,876 x g, 10 minutes, 4 °C). Pellets were resuspended using solubilizing buffer (50 mM Tris, 300 mM NaCl, 10 mM Imidazole, 6 M Urea, pH 8.0), and lysed via sonication (30 seconds on, 30 seconds off) at 50% duty cycle (Branson Sonifer, 15 mm tip) and agitated overnight at 4 °C. Cell lysates were centrifuged (41,657 x g, 4 °C, 1 hour) and supernatants filtered using a 0.45 µM membrane syringe filter. Clarified supernatants were added to 5-10 mL of nickel nitriloacetic acid (Ni-NTA) resin and batch bound (4 °C, 1 hour) prior to purification. The mixture was poured into a glass fritted column and the flow-through fractions collected. Purification would then proceed as follows; the column was washed with 4 column volumes (CVs) of Wash Buffer (50 mM Tris, 300 mM NaCl, 20 mM Imidazole, pH 8.0) followed by elutions using 0.5 CV, 2 CVs of Elution Buffer (50 mM Tris, 300 mM NaCl, 500 mM Imidazole, pH 8.0) collected as two fractions. The 2 CVs elution fraction containing the target protein was dialyzed into 1 L of dialysis buffer (50 mM Tris, 300 mM NaCl, pH 7.5) containing 1 mM EDTA (1 hour room temperature) followed by dialysis against 3L of the dialysis buffer (4 °C, overnight). VL147N was dialyzed using 4 L of dialysis buffer (50 mM Tris, 150 mM NaCl, pH 7.5). All dialysis was performed with Spectra/Por 3 Dialysis Membrane Standard RC Tubing, 3.5 kDa MWCO

### Preparation of Tag-free VHP101C and VL147N

Following the dialysis, Super TEV protease was then added at a 25:1 molar ratio (protein:protease) with DTT (1 mM final concentration). Cleavage reactions were incubated (37 °C for 2-2.5 hours) and gently agitated.

Following cleavage, the mixture was purified by gravity flow IMAC. The mixture was poured over pre-equilibrated resin and the FT collected as a single fraction. Two Wash fractions (2 CVs each) and a single Elution fraction (2 CVs) were collected. The FT and 1^st^ Wash fraction were typically combined and dialyzed into PIPES Buffer (20 mM PIPES, 50 mM NaCl, pH 6.8).

If necessary, protein samples were concentrated in 10 minute increments at 4 °C using a Sorvall Legend X1R centrifuge (ThermoScientific). VL147N was concentrated at 1500-2000 rpm using a Pierce^TM^ Protein Concentrator PES, 10 kDa MWCO, 5-20 mL (ThermoScientific). VHP101C was concentrated at 5000 rpm using a Macrosep Advance Centrifugal Device with 3K MWCO (Pall Corporation).

### Preparation of Tag-free Link-HP (TEV Cleavage and RP-HPLC)

Link-HP’s purification tags were removed via TEV cleavage as described for other constructs and IMAC purified to remove the free tags and His-tagged Super TEV protease. The protein was then concentrated at 1500 rpm with a Pierce^TM^ Protein Concentrator PES, 10 kDa MWCO, 5-20 mL (ThermoScientific).

Link-HP underwent final purification by RP-HPLC. This was achieved on a Dionex Ultimate 3000 HPLC instrument equipped with a Phenomenex Jupiter® 10 μm C4 300 Å LC Column (250 x 10 mm). For these separations, the aqueous phase consisted of 95% H_2_O, 5% MeCN and 0.1% formic acid while the organic phase was 100% MeCN and 0.1% formic acid at a flow rate of 4.0 mL/min. Mobile phase ramps were set as follows: hold at 5% organic for 1.0 min, linear gradient of 5–35% organic from 1.0–10.0 min, linear gradient of 35-90% from 10.0-15.0 min, hold at 90% organic 15.0–17.0 min, linear gradient of 90–10% organic 17.0–17.1 min, re-equilibrate at 5% organic 17.1–20.0 min. Pure Link-HP was collected as it eluted between 11.0 and 12.0 minutes, as detected via UV-Vis.

HPLC fractions of purified Link-HP were then concentrated by rotary evaporation. The flask was immersed into a water bath set to 50 °C and rotated at 50-70 rpm for 20 minutes. This process removed the bulk of the acetonitrile and resulted in a ∼30% decrease in the total volume. The absorbance of the sample was then measured at 280 nm (assuming the protein was dissolved in pure water) to give an estimate of the final protein yield.

Samples obtained from rotary evaporation were flash frozen and lyophilized using a FreeZone 2.5 Liter -50 °C Benchtop Freeze Dryer (LABCONCO) [(0.014 mBar), condensing coil at -51 to -55 °C, 24-48 hours].

Based on the sample absorbance prior to lyophilization, appropriate volumes of PIPES buffer (20 mM PIPES, 50 mM NaCl, pH 6.8) were added to the lyophilized proteins to reach the desired concentration. The solutions were then incubated at room temperature for 5-10 minutes to completely resuspend the purified protein. Isotopically labeled, lyophilized Link-HP was reconstituted with PIPES buffer containing 25 mM PIPES.

### Mass-spectrometry for protein identification

The identity of protein samples was confirmed via electrospray ionization mass spectrometry (ESI-MS) (**Fig. S2B**). This was performed using either an Advion CMS Expression^L^ mass spectrometer or an Agilent 6545XT AdvanceBio LC/QTOF. In the case of the Advion CMS Expression^L^, the mass spectrometer was interfaced with a Dionex Ultimate 3000 HPLC system. Separations upstream of the mass spectrometer were achieved with a Phenomenex Aeris^TM^ 3.6 μm WIDEPORE C4 200 Å column (100 x 2.1 mm) [aqueous (95:5 H_2_O/MeCN, 0.1% formic acid) / organic (MeCN, 0.1% formic acid) mobile phase at 0.3 mL/min, method: hold 10% organic 0.0-1.0 min, linear gradient of 10–90% organic 1.0–7.0 min, hold at 90% organic 7.0–9.0 min, linear gradient of 90–10% organic 9.0–9.1 min, re-equilibrate at 10% organic 9.1–12.0 min]. LC-MS data from this system were analyzed using Advion Data Express software, and protein charge ladders were deconvoluted using Applied Biosystems/SCIEX Analyst software.

For analysis with the Agilent 6545XT AdvanceBio LC/QTOF, the mass spectrometer was interfaced with an Agilent 1290 Infinity II UHPLC system. Separations upstream of the mass spectrometer were achieved with a Phenomenex Aeris^TM^ 3.6 μm WIDEPORE C4 200 Å column (100 x 2.1 mm)) [aqueous (H_2_O, 0.1% formic acid) / organic (MeCN, 0.1% formic acid) mobile phase at 0.3 mL/min, method: hold at 10% organic 0.0-1.0 min, linear gradient of 10–90% organic 1.0–9.0 min, hold at 90% organic 9.0–11.0 min, linear gradient of 90–10% organic 11.0–11.1 min, re-equilibrate at 10% organic 11.1–15.0 min]. LC-MS data from this system were analyzed and protein charge ladders were deconvoluted using Agilent BioConfirm software.

### F-Actin Bundling Assays

All components necessary for the procedure (other than the test proteins) were obtained from an actin binding kit (Cytoskeleton Inc., BK001), resuspended with the appropriate buffers according to the manufacturer and stored on ice. All AtVLN4 protein fragments were dissolved or resuspended in PIPES Buffer (20 mM PIPES, 50 mM NaCl, pH 6.8). α-actinin, G-actin, bovine serum albumin (BSA) and the VLN4 proteins of interest were clarified in a Sorvall MX 150 Micro-Ultracentrifuge (ThermoScientific) at 58,000 rpm (151,807 x g) at 4 °C for 1 hour using a S120-AT3 Fixed Angle Rotor to remove soluble protein aggregates. Immediately following clarification, the supernatants were equilibrated on ice for 30 minutes. The concentrations of most protein solutions were determined using absorbances at 280 nm based on estimated extinction coefficients determined using ExPasy ProtParam.(42, 43) Absorbances for VL147N were determined at 205 nm due to its lack of aromatic residues. Its extinction coefficient was estimated using the Nick Anthis Protein Parameter Calculator.(44) G-actin was reconstructed in the presence of 1X polymerization buffer supplied by the manufacturer (50 mM KCL, 2 mM MgCl_2_, 1 mM ATP, 10 mM Tris at pH 7.5), and incubated at room temperature for 1 hour to polymerize the G-actin. Resuspended actin solutions supplied 11-20 µM G-actin. Following polymerization, F-actin was mixed with the appropriate buffers and AtVLN4 protein fragments and incubated at either 4 °C or room temperature for 30 minutes. The tubes were then centrifuged at 12,600 x g for 15 minutes at the incubation temperature using an Eppendorf Centrifuge 5415 D or Eppendorf Centrifuge 5415 R (for precise temperature control).

After centrifugation, supernatants (50-52 µL) were removed from the tubes and mixed with 4X SDS-PAGE loading dye. Pellets were resuspended using 30 µL of nanopure water and mixed with 4X SDS-PAGE loading dye. The supernatant and pellet samples were subsequently analyzed via SDS-PAGE. For tests at higher KCl concentrations (+100 mM KCl), 3 M KCl and PIPES + 3 M KCl solutions were spiked into the control and trial solutions as appropriate. Due to differences in the final volumes of SDS-PAGE samples derived from pellet versus supernatant fractions, a correction factor was applied to any quantitative intensity data obtained from programs such as ImageJ to account for differences in the amount of material loaded into each gel lane. The correction factor is as follows: Supernatant band intensity *1.675 = Corrected supernatant band intensity. For quantitative analysis, the SDS-PAGE band intensities were quantified with ImageJ and with this correction factor taken into account (Supplementary Table 1). The % actin pelleted was calculated from SDS-PAGE actin pellet and supernatant band intensities as %actin pelleted = 100*(pellet)/(pellet + 1.675*supernatant). Data was analyzed using a Welch’s two sample *t*-test with Holm-Bonferroni correction for multiple comparisons.

### Fluorescent visualization of actin bundling

F-actin polymerization and VL147N fragment production was prepared as described to mimic the buffer conditions utilized to probe F-actin bundling via centrifugation as per the manufacturer’s instructions (14 mM Tris, 0.182 mM CaCl_2_, 45 mM KCL, 1.82 MgCl_2_, and 1 mM ATP). For that, samples of F-actin (5 µM) alone and F-actin (5 µM) mixed with VL147N (7.0 µM) or BSA (4.4 µM) were used as stock solution for microscopy imaging. Actin filaments or bundles were stained with rhodamine phalloidin (Abcam, Cambridge, UK) and directly visualized with an Olympus BX53 fluorescent light microscope equipped with a 40X or 100X (1.3 NA) oil objective. Samples were diluted to a final F-actin concentration of approximately 0.5 µM in 1XPBS (VL147N and BSA in the stock mixtures were diluted proportionately) and mounted on 0.01% poly-L-lysine (Sigma, Rockville, MD) coated slides. All images were collected using cellSens standard software (Evident Scientific).

### Solution NMR data collection and processing

VLN4 NMR samples (Link-HP, VL147N, VHP101C) were made to 0.13 -0.25 mM of the desired protein in 500 μL of NMR Buffer. For VL147N and VHP101C this was 20 mM PIPES, pH 6.8, 50 mM NaCl supplemented with 40 μL D_2_O and NaN_3_ (0.02% final concentration). For Link-HP, the NMR Buffer was 25 mM PIPES, pH 6.8, 50 mM NaCl supplemented with 40 μL D_2_O and NaN_3_ (0.02% final concentration). For probing the effects of potassium ion, KCl at 150 mM was added to Link-HP and VL147N samples. The impact of D_2_O on pH was not corrected. The ^1^H resonance of H_2_O (4.78 ppm at 25 °C) was used as an internal reference for all samples. All NMR spectra were collected on a Bruker Avance III HD NMR spectrometer operating at 11.7 Tesla (1H resonance frequency of 500 MHz) and equipped with a room temperature (^1^H/^13^C/^15^N) TXI Probe (Western Washington University, Bellingham, WA, USA). ^15^N-HSQC spectra were acquired via uniform sampling recording with at least 96 scans (typically 128 scans), 2048 direct and 256 indirect points. Thermal equilibration delay was set at over 1.2 seconds for every recording and sample. ^15^N-HSQC spectra of VLN4 samples were recorded at 3 °C (4 °C for Link-HP), and 25 °C. The acquired NMR data was processed with NMRPipe software (8.9 Rev 2016.307.12.43 32-bit). Spectra were analyzed and visualized via NMRViewJ software (One Moon Scientific Inc.).

## Results

### idCHAR software reveals charge segregation in *Arabidopsis thaliana* villin linkers

Like other IDRs, the linker region of plant villins is a dynamic conformational ensemble consisting of a set of semi-structured or semi-disordered states.(45, 46) The balance between these states affects function and is presumably sensitive to environmental parameters including temperature and ionic strength, non-covalent interactions, or possibly post-translational modifications.(45, 47–49) In the case of villins, data suggest that temperature and ionic strength control IDR function and structure. For example, the structure of villin’s C-terminal headpiece domain and the backbone dynamics of the adjacent acidic part of the IDR linker show segmental sensitivity to temperature.(20, 21) In terms of ionic strength, most IDR linkers in plant villins are all classified by the CIDER algorithm(39, 50) as “Janus sequences” (**Fig. S3A**), suggesting they collapse or extend as a function of salt concentration due to the presence of segregated basic and acidic segments (**Figs. 1B, S4**). In order to further assess charge segregation, we developed a novel software tool (idCHAR) which is able to determine the positions separating two segments of opposite charge within an IDR. The software also quantifies the level of charge segregation to determine whether the charge separation is significant or not.

Application of idCHAR to the linker portions of the five villin variants from *A. thaliana* indicated that four of them (AtVLN2-AtVLN5) display a distinct charge partitioning pattern which includes a ∼150-residue basic (pI 10-11) N-terminus and ∼40-residue acidic (pI 4-5) C-terminus (**Fig S4**). The linkers in AtVLN2-AtVLN5 were also found to have relatively high charge segregation score values (idCHAR score >25). The higher idCHAR score corresponds to a greater excess of positively charged residues (Arg, Lys) over negatively charged residues (Asp, Glu) in the N-terminal segment and a corresponding greater excess of negatively charged residues over positively charged residues in the C-terminal segment. For example, in the 191-residue linker in AtVLN4, idCHAR identified positions 863-864 as producing the highest score (value of 33) as compared with all other positions within the sequence. **Figure 1** illustrates this outcome, showing the basic N-terminal region (pI 11.7) transitioning to the acidic C-terminal region (pI 4.1) at these residues. Contrary to AtVLN2-AtVLN5, the linker in AtVLN1 was found to have no appreciable acidic segment as the idCHAR score was low indicating no comparable charge segregation in the overall basic linker (**Fig. S4**). Intriguingly, idCHAR analysis identified a short distinct 26-residue basic segment (pI 11.8) within the AtVLN1 linker with 9 basic residues and no acidic residues: RKLASLKGKKTSTKRSSGSQYRSQSK. Without this segment, the rest of the linker is only slightly basic (pI 8.1) with basic and acidic residues alternating. At the moment, the role of this highly basic 26-residue segment in AtVLN1 linker remains untested. It is noteworthy that the idCHAR charge segregation scores indicate clustering of the five AtVLN linker sequences into three distinct groups (**Fig. S4**), which is in line with the established perception that plant villins form three phylogenetic groups: AtVLN1, ATVLN2/AtVLN3 and AtVLN4/AtVLN5.(17, 51) Outside of *A. thaliana*, idCHAR analysis of villin proteins in *Zea mays* (corn) also revealed similar groupings and comparable charge partitioning. Specifically, linkers in the four ZmVLN proteins analogous to AtVLN2-AtVLN5 possessed similar basic (∼150 amino acid residues) and acidic (40-50 amino acid residues) segments, while ZmVLN1 possessed no significant charge partitioning (data not shown).

### The basic fragment of the AtVLN4 linker (VL147N) bundles F-actin

In considering the relevance of charge segregation, we noted that charge patterns in plant villin linkers were analogous to those observed in the 830-residue IDR of supervillin. In supervillin, basic segments within its IDR are central for F-actin binding and bundling, which led us to hypothesize that basic regions within villin linkers may have similar functions.(52–54) To test this, we chose to probe the linker from AtVLN4 because this variant has the highest reported affinity for F-actin (0.25 µM) among the five plant villins (17) and produced the highest idCHAR score in the linker analysis described above. This linker was also classified by the CIDER algorithm as a “Janus sequence”, suggesting that if F-actin bundling was observed this property may be dependent on environmental conditions such as ionic strength or temperature.(39, 40)

Initially, a construct consisting of the 147-residue basic region of the AtVLN4 linker and three residues added to the C-terminus to minimize possible C-terminal proteolysis was tested for actin bundling. This fragment, referred to as VL147N (<u>V</u>illin <u>L</u>inker, <u>147</u> <u>N</u>-terminal residues, **Fig. 1**), was combined with F-actin and subjected to low-speed centrifugation following by SDS-PAGE analysis. VL147N was assayed at three concentrations (1.6 μM, 3.2 μM and 5.9 μM) at a temperature of 21 °C as per established procedures.(55, 56) Under these conditions, we observed that the bulk of F-actin was reproducibly found in the pellet fractions, suggesting that VL147N was capable of actin bundling (**Fig. 2A-B**, **Figs. S5-S7**). These results mirrored those of a positive control using the known F-actin bundling protein α-actinin and were markedly different from a negative control using BSA, which showed little to no actin in the pellet fraction (**Fig. 2C**, **Fig. S8**). In isolation, F-actin and VL147N generally did not pellet, though some cases insignificant self-pelleting of F-actin was noted, which is in line with the actin supplier levels (for examples see **Figs. S8A**, right panel).

**Figure 2.**
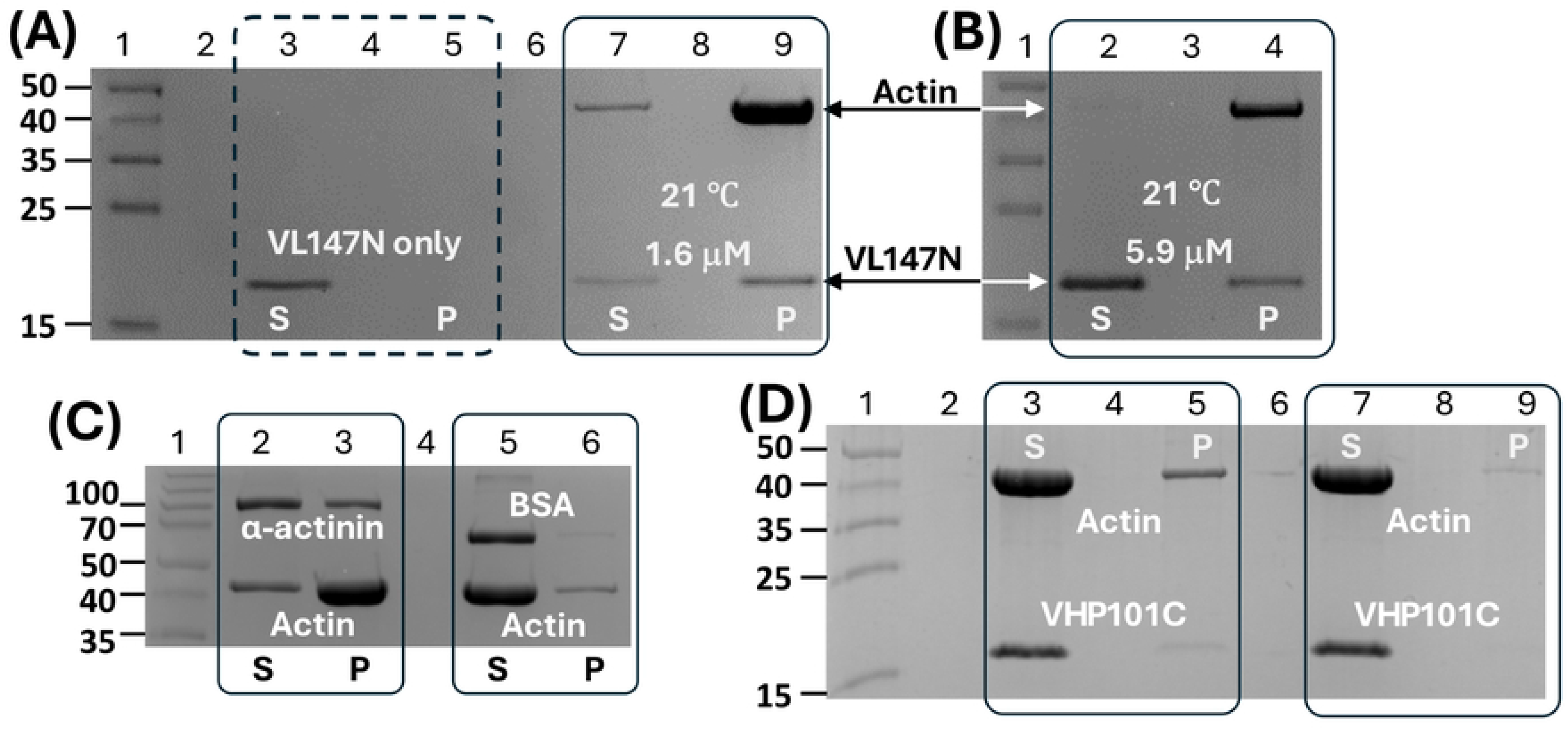
VL147N bundles F-actin at 21 °C. **(A)** VL147N at 1.6 μM pulls down F-actin under low-speed centrifugation conditions. Lane 1: molecular weight standards (in kDa); lanes 3 and 5 in a dashed frame: centrifugation supernatant (S) and pellet (P) of VL147N in isolation (1.6 μM); lanes 7 and 9: centrifugation supernatant (S) and pellet (P) of VL147N (1.6 μM) mixed with F-actin. Solid rectangular frames in this and other panels indicate the supernatant (S) and pellet (P) fractions of the same pull-down experiment. **(B)** VL147N at 5.9 μM pulls down F-actin. Lane 1: molecular weight standards (in kDa); lanes 2 and 4: centrifugation supernatant (S) and pellet (P) of VL147N (5.9 μM) mixed with F-actin. **(C)** Control experiments at 21 °C. Lane 1: molecular weight standards (in kDa); lanes 2 and 3 (positive control): centrifugation supernatant (S) and pellet (P) of α-actinin (∼100 kDa; 1.0 μM) mixed with F-actin (10 μM); lanes 5 and 6 (negative control): centrifugation supernatant (S) and pellet (P) of BSA (∼66 kDa; 1.6 μM) mixed with F-actin (10 μM). **(D)** VHP101C at 5.9 μM does not pellet F-actin at 21 °C. Lane 1: molecular weight standards (in kDa). Lanes 3 and 5: centrifugation supernatant (S) and pellet (P) of VHP101C (5.9 μM) mixed with F-actin. Lanes 7 and 9 represent a duplicate trial of the same experiment.

In these co-sedimentation experiments at 21 °C, the fraction of F-actin remaining in the supernatant appeared to depend on the concentration of VL147N in the mixture. For VL147N at 1.6 μM, a noticeable actin band in the supernatant fraction was present for every trial (**Fig. 2A**, **Fig. S7**), whereas little to no actin supernatant bands were observed when VL147N was present at 5.9 μM (**Fig. 2B**, **Fig. S5)**. The pelleting experiment performed with VL147N at 3.2 μM demonstrated similar outcomes to VL147N at 1.6 μM with a noticeable actin band in the supernatant fractions (**Figs. S6A, S6C**). However, at all VL147N concentrations tested with F-actin, the bulk of actin was consistently found in the pellet fractions.

In addition to VL147N, we also assessed actin bundling in the presence of other AtVLN4 linker fragments. One of them, VHP101C, represents the rest of the AtVLN4 C-terminus and its 101 residues include the acidic portion of the IDR linker fused with the headpiece domain (**Fig. 1**). In contrast to VL147N, VHP101C exhibited essentially no actin pull-down at 21 °C, suggesting a lack of actin bundling capability in the section of AtVLN4 C-terminal to VL147N (**Fig. 2D**).

Having observed actin co-sedimentation using centrifugation, we next verified the actin bundling activity of VL147N through direct visualization of actin filaments using fluorescent microscopy. As shown in **Fig. 3**, clear bundles were observed in samples containing VL147N and F-actin, whereas these were absent in samples of F-actin alone or F-actin in combination with BSA. Overall, this provided an independent confirmation of the ability of VL147N to initiate actin bundling.

**Figure 3.**
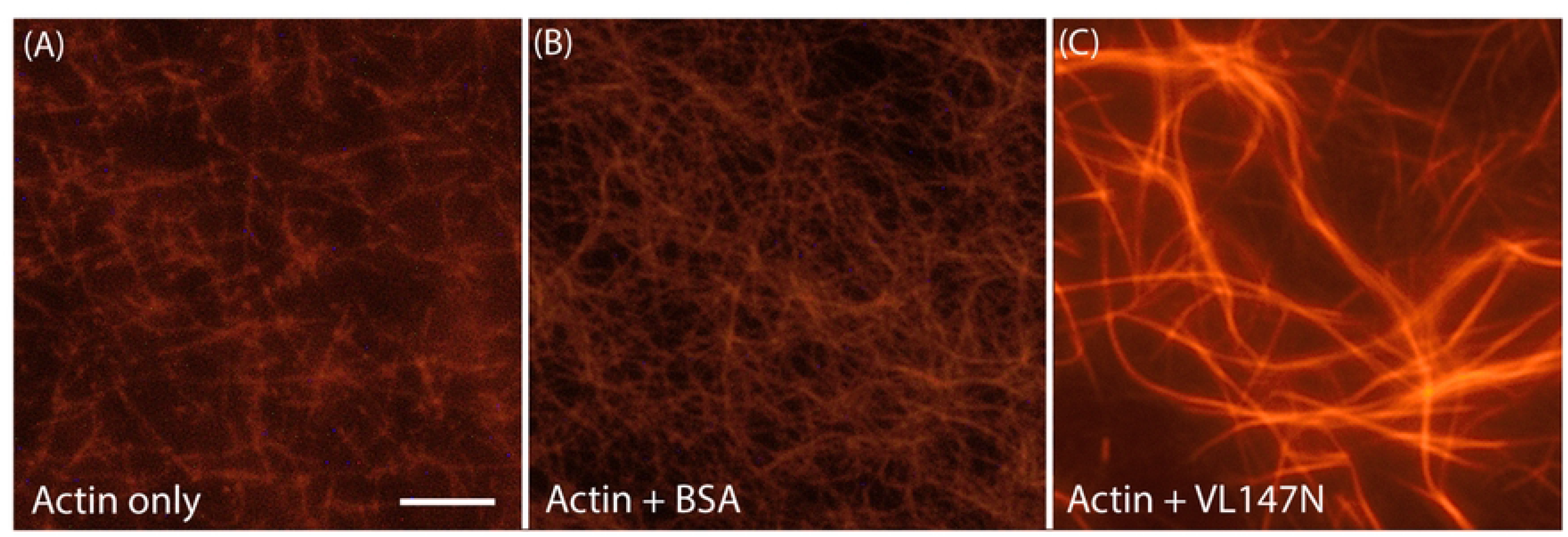
Visualization of F-actin bundling by VL147N. Fluorescent microscopy images of F-actin stained with rhodamine-phalloidin. **(A)**: F-actin alone; **(B):** F-actin mixed with BSA; **(C)**: F-actin combined with VL147N, indicating that actin bundles are formed only in the presence of VL147N. Scale bar = 10 μm.

### VL147N bundles F-actin at 21 °C more effectively than at 4 °C

Having observed that VL147N is capable of bundling F-actin, we began to explore whether this property was sensitive to temperature. To this end, F-actin bundling by VL147N was tested at 4 °C (**Figs. 4A-D**) to complement the 21 °C trials described above. Notably, both 4 °C and 21 °C represent physiologically relevant environmental conditions for plants. Pelleting of F-actin by VL147N was demonstrated at both temperatures for VL147N concentrations in the range of 1.6 μM to 5.9 μM. At 4 °C, less than 40% of actin was found in the pellet fractions when VL147N was present at 1.6 μM (**Fig. 4B**, **Figs. S7B**, **S7D** and **Fig. 5A**; **Table S1**). In contrast, more than 80% actin was pelleted at 21 °C (**Fig. 4A**, **Fig. S7A**, **S7C** and **Fig. 5**; **Table S1**). The effect of significant reduction of the pelleted F-actin fraction was also observed when F-actin was mixed with VL147N at 3.2 μM (**Figs. S6A**, **S6C** vs. **Figs. S6B**, **S6D**) and at somewhat lower scale but still consistently when VL147N was present at 5.9 μM (**Fig. 4C** for 21 °C vs. **Fig 4D** for 4 °C; **Figs. S5A**, **S5C** vs. **S5B**, **S5D**).

**Figure 4.**
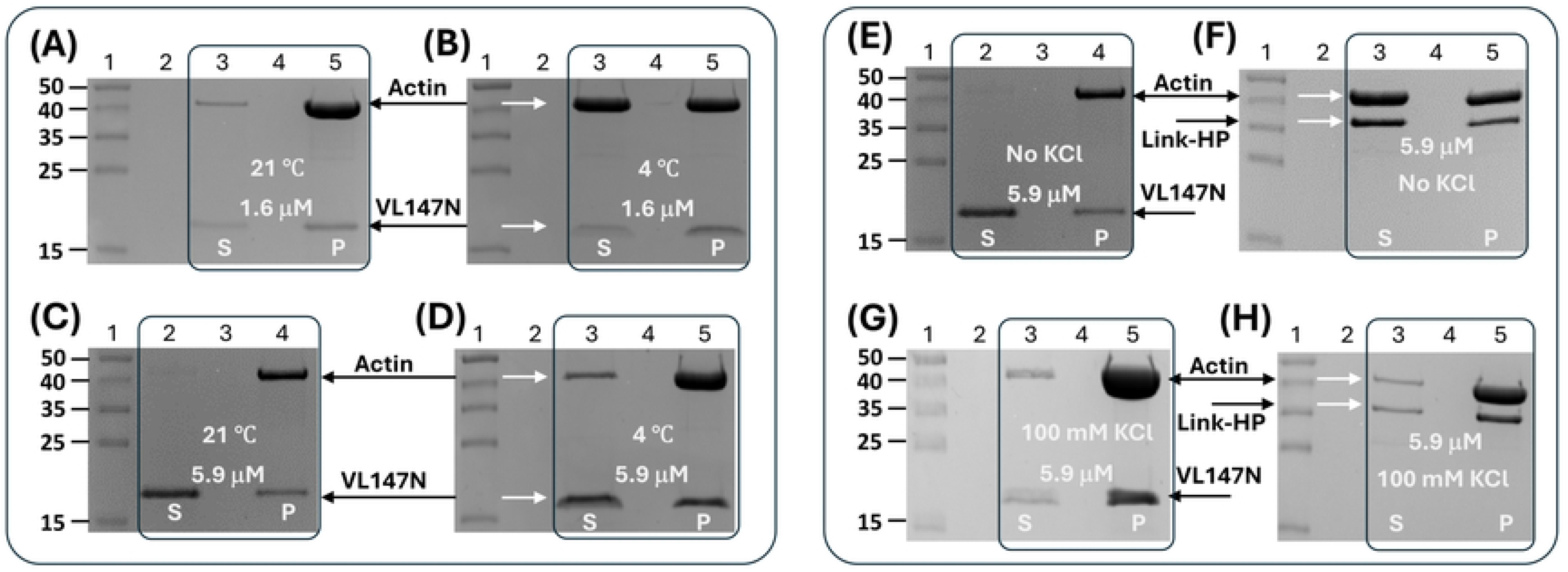
(A-D) VL147 bundles F-actin at 21 °C stronger than at 4 °C. Black lines and white arrows mark the positions for the actin and VL147N gel bands. **(A)** Lane 1: molecular weight standards (in kDa); lanes 3 and 5: VL147N (1.6 μM) mixed with F-actin centrifugation, supernatant (S) and pellet (P) fractions at 21 °C. **(B)** Lane 1: molecular weight standards (in kDa); lanes 3 and 5: centrifugation supernatant (S) and pellet (P) of VL147N (1.6 μM) mixed with F-actin at 4 °C. **(C)** Lane 1: molecular weight standards (in kDa); lanes 2 and 4: centrifugation supernatant (S) and pellet (P) of VL147N (5.9 μM) mixed with F-actin at 21 °C. **(D)** Lane 1: molecular weight standards (in kDa); lanes 3 and 5: centrifugation supernatant (S) and pellet (P) of VL147N (5.9 μM) mixed with F-actin at 4 °C. **(E-H) Link-HP bundles F-actin weaker than VL147N but the bundling is restored by 100 mM KCl.** All the tests here were done at 21 °C and in the presence of 50 mM NaCl. Concentration values of 5.9 μM were used for VL147N and Link-HP. **(E)** Lane 1: molecular weight standards (in kDa); lanes 2 and 4: centrifugation supernatant (S) and pellet (P) of VL147N (5.9 μM) mixed with F-actin in the absence of KCl at 21 °C. **(F)** Lane 1: molecular weight standards (in kDa); lanes 3 and 5: centrifugation supernatant (S) and pellet (P) of Link-HP (5.9 μM) mixed with F-actin in the absence of KCl at 21 °C. **(G)** Lane 1: molecular weight standards (in kDa); lanes 3 and 5: centrifugation supernatant (S) and pellet (P) of VL147N (5.9 μM) mixed with F-actin in the presence of 100 mM KCl at 21 °C. **(H)** Lane 1: molecular weight standards (in kDa); lanes 3 and 5: centrifugation supernatant (S) and pellet (P) of Link-HP (5.9 μM) mixed with F-actin in the presence of 100 mM KCl at 21 °C.

**Figure 5.**
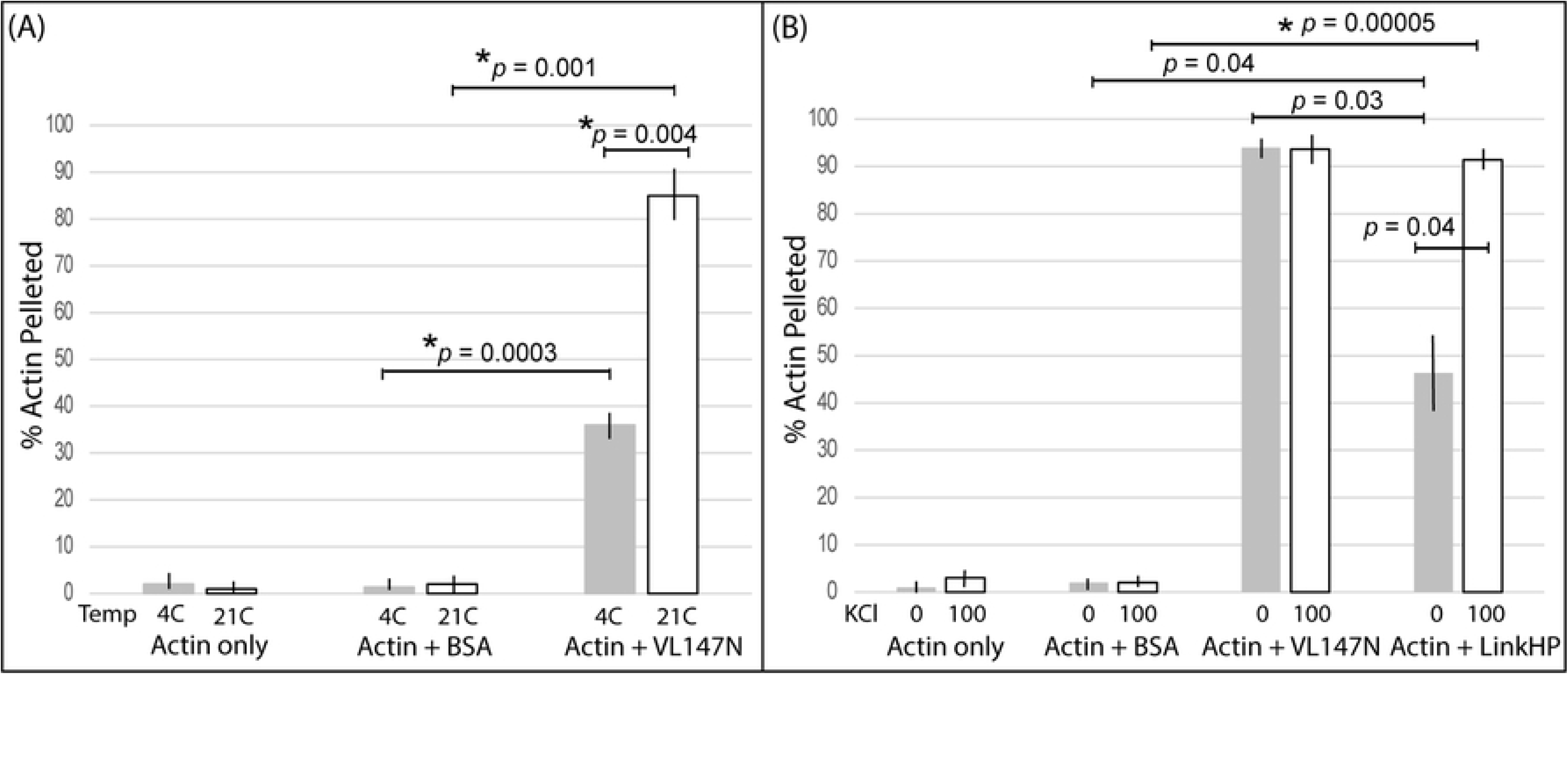
F-actin bundling by linker fragments of AtVLN4 is responsive to temperature and KCl. Also see **Table S1**. Bars indicate the mean of three replicates ± SE. *p*-values are indicated over horizontal bars for selected comparisons; actin-only comparisons were similar to Actin +BSA comparisons and were omitted for clarity. Statistics were calculated using Welch’s two-sample *t*-test with a Holm-Bonferroni correction to ensure an experiment wide α = 0.05. Asterisks indicate significance after correction for multiple comparisons. **(A)** F-actin bundling capacity of VL147N was significantly reduced at low temperature (4 °C) compared to 21 °C. Experiments were performed in the absence of KCl, VL147N concentration was 1.6 µM; BSA concentration was 4.4 µM for all replicates (n = 3). **(B)** Link-HP F-actin bundling increased in the presence of 100 µM KCl. Experiments were performed at 21 °C, VL147N concentration was 5.9 µM; Link-HP concentration was 5.9 µM, BSA concentration was 4.4 µM for all replicates (n = 3). VHP101C displayed limited capacity for actin bundling in our tests (not shown).

### AtVLN4 linker fragments respond differently to the presence of KCl

Our AtVLN4 254-residue Link-HP fragment represents the entire linker, including the VL147N, fused with the headpiece domain (Fig. 1A). Under the same conditions as VL147N, the fraction of actin in the supernatant for Link-HP is noticeably higher than the supernatant fraction for VL147N, indicating weaker F-actin bundling by Link-HP (**Fig. 4E** vs. **Fig. 4F**; **Figs. S9A**, **S9B**). This weaker bundling by Link-HP compared to VL147N was also observed for the lower value of concentrations tested for the two AtVLN4 fragments (1.6 μM) (**Figs. S9C**, **S9D**). Thus, while Link-HP contained the full VL147N sequence, the presence of the headpiece domain and the acidic segment of the linker appeared to impede actin bundling.

As a possible explanation for the lower bundling ability of Link-HP, we considered the fact that the full length villin 4 linker present in this construct possesses clear regions of charge segregation (**Fig. 1**). In solution, regions of opposing charge may associate in an intra- or intermolecular fashion, leading to occlusion of the actin binding/bundling sites in the VL147N segment and a reduction in actin bundling by Link-HP. We therefore hypothesized that the actin bundling ability of Link-HP might be enhanced by increasing ionic strength, which could mask any inhibitory electrostatic interactions. To test this, we repeated our actin bundling assays in the presence of KCl, which is a physiologically relevant salt for plants. Indeed, introduction of KCl at 100 mM increased the capacity of Link-HP to bundle F-actin to a similar level as for isolated VL147N (**Fig. 4G** vs. **Fig. 4H**, **Fig. 4F** vs. **Fig. 4H**, **Fig. S10**, **Fig. 5**, **Table S1**) and for both fragments the intensity of the supernatant actin band was minimal. Importantly, the capacity of VL147N to bundle F-actin was not affected by the presence or absence of KCl (**Fig. 4E** vs. **Fig. 4G**, **Fig. 5B**, **Table S1; Fig. S11**).

In addition to VL147N and Link-HP, the VHP101C fragment (**Fig. 1**) described above was tested under the same sets of conditions. VHP101C did not to pellet F-actin at 21 °C or 4 °C at any significant level (**Fig. 2D** and **Fig. S12A** respectively). The amount of actin pelleted by VHP101C at 100 mM KCl (**Fig. S12B**) was low and on par with that of actin pelleted in isolation (**Fig. S13A**) or by BSA (negative control, **Fig. 2C, Fig. S13B**).

Taken together, these findings are consistent with a recent report indicating that the full AtVLN5 and AtVLN4 linkers are able to bind F-actin.(22) Further, our verification that VL147N and Link-HP, but not VH101C, have actin bundling capacity is consistent with Zhuang et al., (22) who determined that a subsection of the basic portion of the closely related AtVLN5 linker was critical for actin bundling by the core-linker fusion. Our observation that the headpiece-containing Link-HP has a reduced F-actin bundling capabilities also is in agreement with the observations of these authors who determined that the headpiece is not needed for F-actin bundling by AtVLN5.

Quantitative SDS-PAGE band intensity analysis supported our qualitative conclusions indicating that the bundling capacity of certain linker fragments of AtVLN4 is altered by environmental conditions. The percent actin pelleted calculated through gel band intensities using ImageJ are summarized in **Fig. 5** and **Table S1**. High actin bundling values, greater than 75% with average values in 85-94% range, indicate strong pelleting (bundling) by the VL147N and Link-HP (in high K^+^), but not for VHP101C (**Table S1**). Two experimental treatments relevant for plant physiology, temperature and ionic strength, resulted in changes in F-actin bundling, with average values between 36-47%. Specifically, VL147N pellets (bundles) significantly less F-actin at 4 °C than at 21°C (Fig. 5A). Similarly, we determined that Link-HP bundles F-actin weakly at marginally significant levels in the absence of KCl compared to in the presence of 100 mM KCl (Fig. 5B). While the observed trend-level evidence for Link-HP and KCl did not reach statistical significance after appropriate multiple comparison correction, future studies with a larger sample size and KCl concentrations clearly warrant investigation.

### Link-HP contains the majority of NMR spectral features of the headpiece and VL147N

We next employed solution NMR spectra ({^1^H-^15^N}-HSQC) as a general probe for the conformational states of our villin fragment samples. **Fig. 6** shows overlays of the NMR spectra of Link-HP with the headpiece and VL147N (**Fig. 6A** and **Fig. 6B**, respectively). The VL147N spectrum recorded at 3 °C exhibited backbone amide signals of varying intensity with a total peak number of 130 signals if the weakest signals are discounted. This corresponded to nearly 100% coverage of the ∼150-residue fragment with 21 prolines. For the majority of the headpiece and VL147N signals, there was a closely matching signal from Link-HP (**Fig. 6**). This observation suggests that the backbone conformational ensembles within Link-HP are modular: the conformational state of the headpiece within Link-HP and in isolation are very similar and the same is true for VL147N.

**Figure 6.**
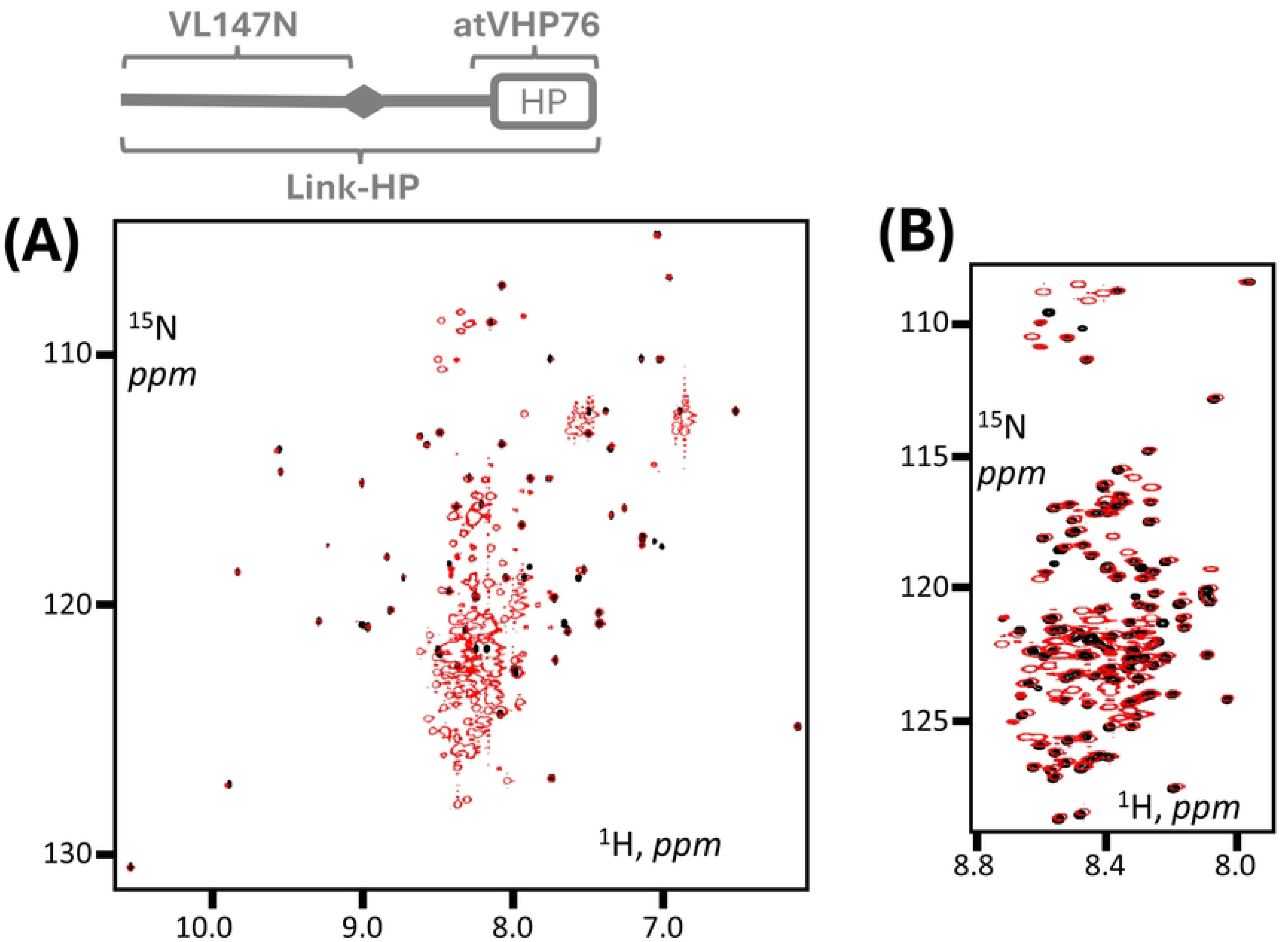
{^1^H-^15^N}-HSQC spectra of Link-HP, headpiece and VL147N (11.7 Tesla, 500 MHz, ^1^H frequency). **(A)** Link-HP (red) and headpiece (atVHP76 (20), black) at 25 °C; **(B)** Link-HP (4 °C, red) and VL147N (3 °C, black). The vast majority of the headpiece and VL147N signals have a counterpart in the respective Link-HP spectrum, which indicates structural modularity for Link-HP.

### Link-HP and VL147N NMR spectra indicate local structural differences in the absence of KCl

In addition to the observation of significant spectral similarities between Link-HP and VL147N, we also observed a certain number of cases (at least six) when an NMR signal emanating from a VL147N residue had no match in the Link-HP spectrum (**Fig. 7**). Here we compared NMR spectra at low temperature (3-4 °C) as these data have greater sensitivity than those at 25 °C, with significantly more peaks observed for both samples. These observations indicated that the local chemical environment and conformations adopted by these several residues in VL147N, as well as their backbone dynamics, differ from those in Link-HP.

**Figure 7.**
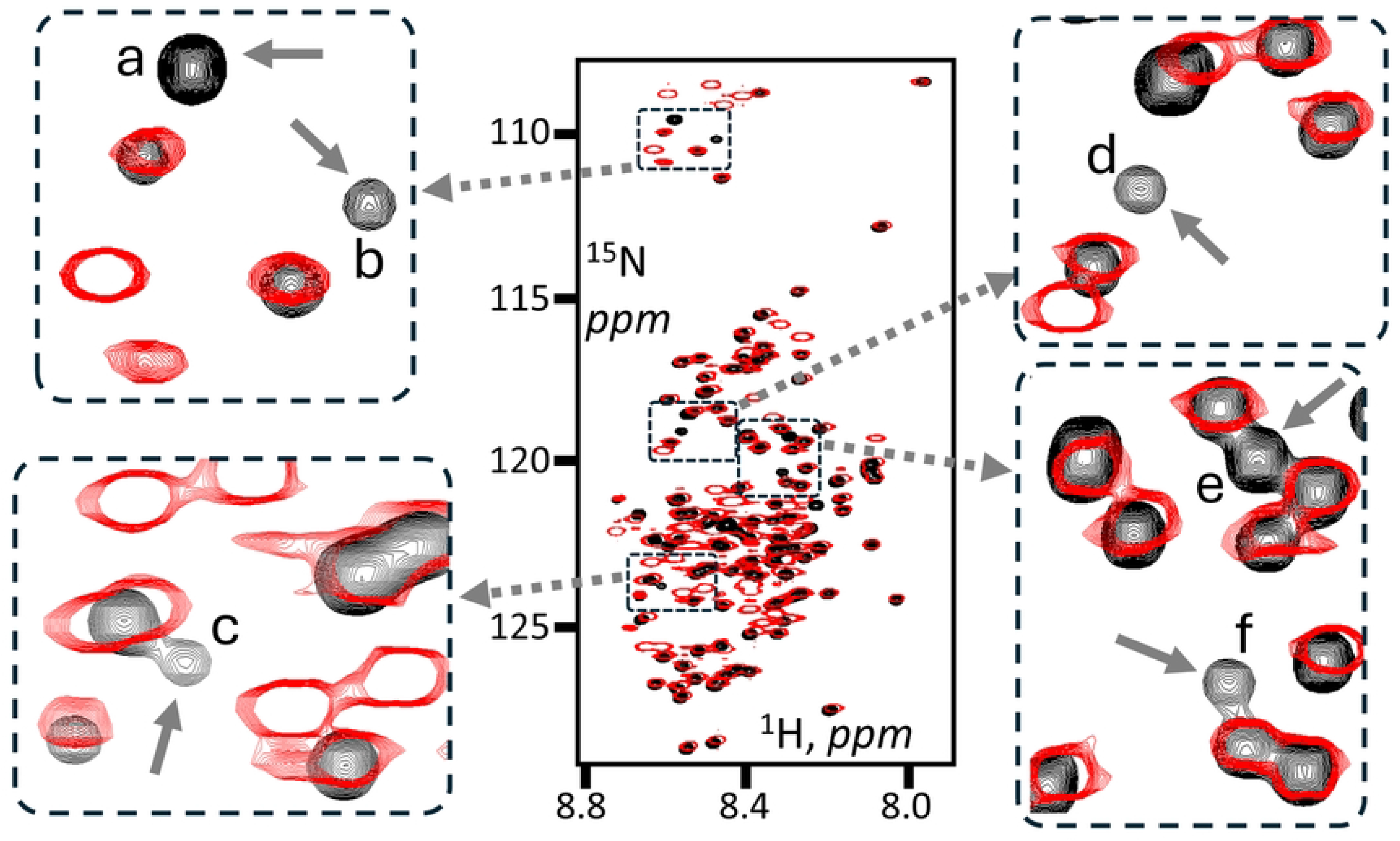
{^1^H-^15^N}-HSQC spectral differences between Link-HP and VL147N (11.7 Tesla, 500 MHz, ^1^H frequency). Solid arrows point to at least six cases of VL147N signals (3 °C, black) which have no match in the Link-HP data (4 °C, red). These signals are also labeled with letters, a-f. The buffers in both cases lacked KCl to mimic the environment of the F-actin bundling tests described in Figs. 2 and 3.

### High KCl restores NMR spectral similarity between Link-HP and VL147N

As described above, high KCl levels restored the capacity of Link-HP to pellet F-actin (**Fig. 4**). This might be explained by the effect of K^+^ ions on the structure or fold of Link-HP. To assess this, {^1^H-^15^N}-HSQC NMR spectra were recorded at an elevated KCl concentration to check whether KCl restores structural similarity between VL147N and the respective segment within Link-HP (**Fig. 8**). This comparison revealed that KCl at 150 mM restores the presence of four out of six signals which were missing in the Link-HP spectrum recorded in the absence KCl (**Fig. 8**). A possible fifth missing signal may have also reappeared in the Link-HP spectrum in high KCl, although it was more distant from its possible counterpart (**Fig. 8**)

**Figure 8.**
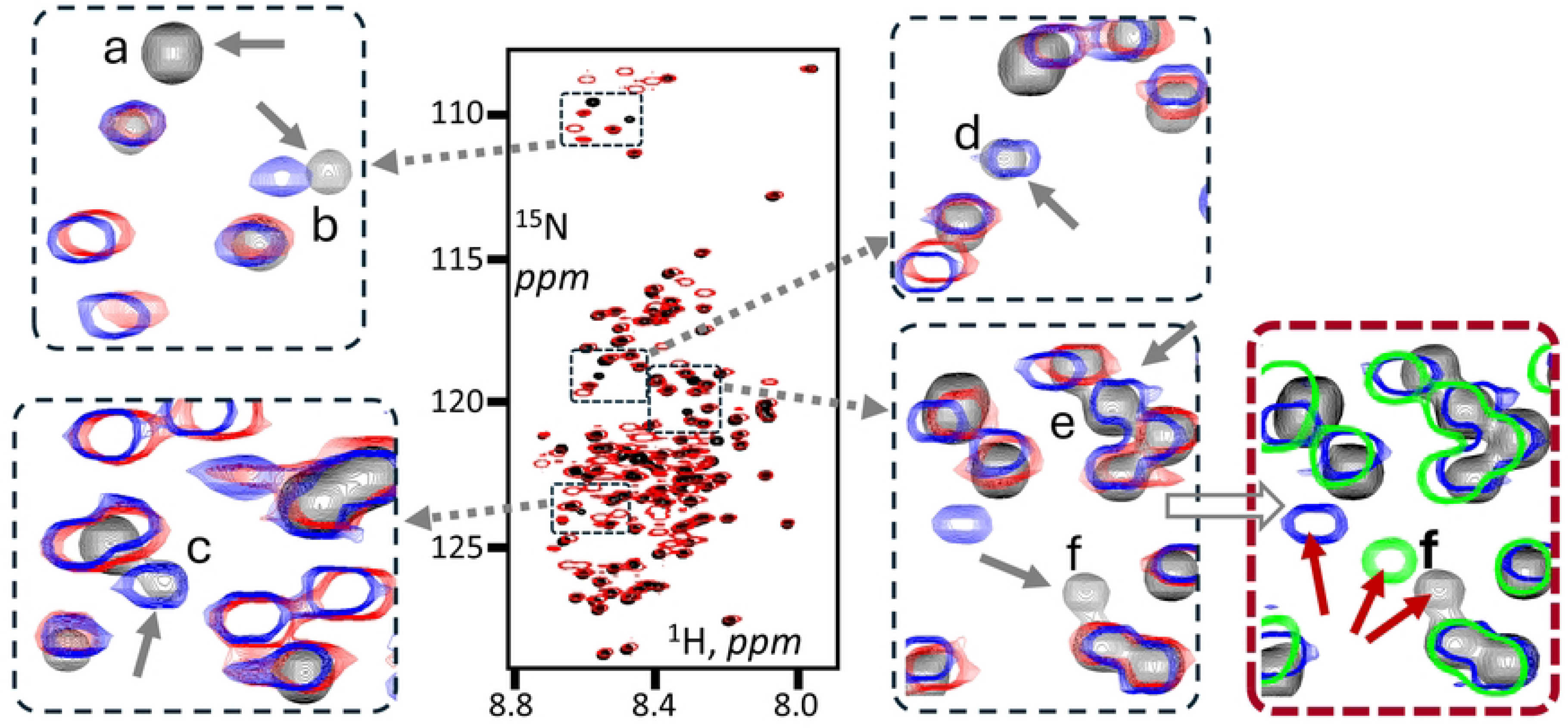
Spectral {^1^H-^15^N}-HSQC differences between Link-HP with and without KCl as compared with VL147N. The data was recorded at 11.7 Tesla (500 MHz, ^1^H frequency), the color-coding shown is based on the scheme used in Fig. 7: VL147N without KCl (3 °C, black), Link-HP without KCl (4 °C, red), and Link-HP with 150 mM KCl (4 °C, blue). The solid arrows point to at least six cases of VL147N signals which have no match in the KCl-free Link-HP data. These signals are also labeled with letters, a-f. Out of these six signals, at least four (b, c, d, e) are restored in Link-HP once KCl (150 mM) was added. The fifth peak “f” is shown in more detail in the inset on the right (red frame): the signals in green represent VL147N in 150 mM KCl.

Unlike for Link-HP, a high level of KCl did not perturb the VL147N NMR spectrum in a significant way. With KCl added, the spectra retained the same number of signals with only minor chemical shift variations (≤0.03 ppm and <0.1 ppm in the ^1^H and ^15^N dimensions, respectively) for data recorded at 3 °C (**Suppl. Fig. 14**) or 25 °C (**Suppl. Fig. 15**). Even the most notable KCl-induced spectral alterations were minor. One such shift is shown in **Fig. 8** (signal “f” in the red frame inset: black vs. green signals). Moreover, the effects of KCl on the spectra of Link-HP and VL147N at 25 °C are similar to those at 4 °C (**Suppl. Figs. 16, 17**).

## Discussion

Together with other key proteins,(57, 58) villins play significant roles in the regulation of the actin cytoskeleton, which impacts a variety of plant processes. (12, 22, 59–62) Here, we have described a new software tool (idCHAR) for assessing charge separation in IDR sequences and used it to analyze five villin isoforms from *A. thaliana*. Based on the charge segregation revealed by idCHAR, we also investigated the actin bundling ability of the 191-residue AtVLN4 linker using full-length (Link-HP) and truncated N-and C-terminal fragments (VL147N and VHL101C) (**Fig. 1**). Our functional analysis demonstrated that the N-terminal basic segment (VL147N) of the AtVLN4 linker is able to bundle F-actin, which was confirmed through actin co-sedimentation and fluorescence microscopy. This bundling ability was sensitive to temperature, with significantly less bundling observed at lower temperature. Our data also suggests that F-actin bundling using our large linker fragment (Link-HP) was impacted by the presence of KCl, which was not a factor that influenced actin bundling for VL147N (Fig. 5). Notably, the C-terminal fragment VHP101C was shown not to bundle F-actin under any temperature and K^+^ conditions tested. While the improved bundling in the presence of KCl was not statistically significant (p < 0.05, Fig. 5B) after our stringent multiple comparison correction, our data provide clear trend-level evidence based on ionic strength responsiveness for the Link-HP fragment that could be remedied by a larger sample size. This quantitative data combined with the potential for intra-or intermolecular interference based on charge separations of Link-HP suggests a potential role of ionic strength in VLN linker bundling ability and the need for a more comprehensive investigation in the future.

In addition to functional assays, solution NMR analysis suggests that the VL147N segment within Link-HP adopts a similar conformation to VL147N in isolation (**Fig. 6**). However, certain NMR chemical shift differences and signal disappearances indicate different local conformation(s), alternative dynamics regimes or different non-covalent interfaces for VL147N in isolation versus within Link-HP (**Fig. 7**). The addition of KCl to Link-HP helps to restore the structural and/or dynamics properties of the VL147N segment within Link-HP to the states observed in the spectra of isolated VL147N (**Fig. 8**), whereas KCl does not perturb isolated VL147N in a significant way (**Suppl. Figs. 12-13**, **Fig. 8**).

Due to the high sequence homology between AtVLN4 and the basic segments of linkers in related proteins in other plants (**Suppl. Figs. 1B, 1C**) as well as within the headpiece domains,(20) we speculate that the ability to initiate F-actin bundling is shared by all AtVLN4-like plant villins. While not pursued here, a more detailed molecular investigation of F-actin bundling by these types of linker sequences would be a valuable future direction, along with possible *in vivo* studies and site-directed mutagenesis to identify residues in villin IDRs that are critical for interactions with actin or responsiveness to environmental changes.

More broadly, our results offer an example of a sizable basic IDR capable of F-actin bundling, which can be compared with a similar example in the related vertebrate protein supervillin (**Suppl. Fig. 1A**). While supervillin’s massive IDR has a ∼340-residue segment capable of F-actin bundling (52, 54), the 147-residue VL147N segment presented here is far shorter and thus may serve as a model to answer basic questions about this highly versatile protein: what is the minimum length for a basic IDR fragment that retains the capacity to bind or bundle F-actin? How is VLN-directed actin bundling impacted by environmental changes? Notably, the 2D {^1^H-^15^N}-HSQC spectra presented here offer strong sensitivity and resolution, suggesting that a combination of low-velocity pull-down assays, fluorescent microscopy and solution NMR may offer a powerful tool kit for these types of investigations. NMR may potentially also allow identification of the specific residues undergoing the K^+^-sensitive conformational or interface changes observed between VL147N and Link-HP. Ultimately, we envision that future studies of this type involving all AtVLN proteins (AtVLN1-5) will shed light on how this gene family may work independently and cooperatively to regulate plant growth and development via actin modifications under changing environmental conditions.

## Supporting Information Captions

**S1 Fig 1**. (**A**): Domains and IDR linkers in plant and animals: the terminal folded core and headpiece (HP) domains are highly homologous whereas the internal IDR linkers are markedly different (short in animals and much larger in plants). The basic parts of the linker are shown as blue lines whereas red lines represent the acidic parts. (**B**) ClustalW Amino acid sequence alignment of the basic region of representative monocot and dicot VLN4 IDRs. Identical amino acids are highlighted in green and asterisks. Similar amino acids are highlighted in yellow and colon. Sequences were aligned using Clustal Omega. Sequence accessions are Glycine max (XP_040867312), Arabidopsis thaliana (NM_001203940), Oryza sativa (NP_001406776), and Zea mays (XP_020400860). (**C**) The amino acid comparison table for the alignment in (**B**).

**S2 Fig**. **(A)** Sequences of VLN4 constructs used in this study. Sequences shown represent proteins following TEV cleavage to remove N-terminal affinity tags. In VLN147N, a tripeptide GRA follows the C-terminal proline to prevent proline-related exoproteolysis. **(B)** Deconvoluted ESI-MS spectra of purified AtVLN4 constructs (calculated MW for VL147N = 16135 Da, calculated MW for VHP101C = 11983 Da, calculated MW for Link-HP = 28031 Da).

**S3 Fig**. **(A) CIDER analysis of the linkers from six plant villin variants, AtVLN1-AtVLN5**. Two AtVLN1 sub-variants were included here: AtVLN 1.1 and AtVLN 1.3, labeled as 1.1 and 1.3, respectively. Variants AtVLN2-AtVLN5 are labeled with a single digit: 2, 3, 4 and 5, respectively. The sequences of the linkers are shown in **Fig. S4** (AtVLN 1.3 as the only one for AtVLN1). The graph shows one dot for every linker sequence with the coordinates f+ and f-representing fractions of positively and negatively charged residues as calculated by the CIDER algorithm. The space between two dashed lines corresponds to the “Janus sequences”: collapsed or extended (context dependent). All the *A. thaliana* villin linkers except that from VLN3 are classified by CIDER as Janus sequences. (**B**) **An example of idCHAR score calculation algorithm** for a model sequence G_1_R_2_D_3_A_4_K_5_P_6_P_7_E_8_L_9_K_10_D_11_A_12_. The charge segregation score is calculated for every sequence position starting from 2 through N-1 (N is the sequence length, 12 in this case) as follows: a) the difference between the number of basic (R, K) and acidic (D, E) residues is calculated for the residues N-terminal to the sampled position, b) the same value for residues C-terminal to the sampled position and c) the score: the difference of the quantities obtained in a) and b). The table shown reports the scores calculated for positions 2 through 11. The residues reporting the highest absolute value of the score are reported as the boundary between the charge-segregated segments (positions 6, 7 here with the score of 2). The positive value of the boundary score indicates a basic N-terminus (GRDAK here) and acidic C-terminus (EAKDA here), the negative score values indicate the opposite scenario, e.g., in sequence G_1_D_2_R_3_A_4_E_5_P_6_P_7_R_8_L_9_D_10_K_11_A_12_, panel **(C)**.

**S4 Fig**. Amino acid sequences of the linkers in AtVLN1-AtVLN5. Charge segregation as identified by idCHAR in the linkers for AtVLN2, AtVLN3, AtVLN4 and AtVLN5 is color-coded: blue for the basic N-terminus and red for the acidic C-terminus. “Boundary” (in black underlined) refers to a short segment between the basic and acidic segments for AtVLN2-AtVLN5. In bold italics is the basic, 26-residue segment (pI 11.8) in AtVLN1 which conveys the bulk of the basic nature for this linker (see main text). The boundary between the basic and acidic segments in AtVLN1 is formally reported by idCHAR but the “acidic” segment in this IDR is only six residues in size with only two acidic ones. In AtVLN4, residues in italics denote the predicted regulatory PEST motif.

**S5 Fig**. F-actin pelleting tests by VL147N at 5.9 µM. **(A)** Pelleting test for VL147N in isolation (lanes 2, 4) and VL147N mixed with F-actin (lanes 6, 8) at 21 °C and VL147N concentration of 5.9 µM, trial 1. **(B)** Pelleting test of VL147N at 5.9 µM in isolation (lanes 3, 5) and trial 1 of F-actin pelleting by VL147N at 5.9 µM, both tests at 4 °C (lanes 7, 9). **(C)** Pelleting tests for VL147N at 5.9 µM mixed with F-actin at 21 °C, trials 2 and 3. **(D)** Pelleting tests for VL147N at 5.9 µM mixed with F-actin at 4 °C, trials 2 and 3. Centrifugation fraction labels on top of the gels are as follows: S, supernatant and P, pellet. The molecular weight (MW) marker values are in kDa. Each round-corner frame encapsulates the supernatant and pellet fractions of the same VL147N + F-actin trial.

**S6 Fig**. Pelleting of F-actin by VL147N at 3.2 µM, temperature probed: 21 °C and 4 °C. Tests were done in triplicates. For 21 °C, panel **(A)** shows trial 1, **(C)** -trials 2, 3; For 4 °C, panel **(B)** shows trial 1, **(D)**-trials 2, 3. Centrifugation fraction labels on top of the gels are as follows: S, supernatant and P, pellet. The molecular weight (MW) marker values are in kDa. The outcomes of all the controls are the same as for the VL147N tests at 5.9 µM (Fig. 2 and Suppl. Fig. 8).

**S7 Fig**. Pelleting of F-actin by VL147N at 1.6 µM, temperature probed: 21 °C and 4 °C. Tests were done in triplicates. For 21 °C, panel **(A)** shows trial 1, **(C)** -trials 2, 3; For 4 °C, panel **(B)** shows trial 1, **(D)**-trials 2, 3. Centrifugation fraction labels on top of the gels are as follows: S, supernatant and P, pellet. The molecular weight (MW) marker values are in kDa. The outcomes of all the controls are the same as for the VL147N tests at 5.9 µM (Fig. 2 and Suppl. Fig. 8).

**S8 Fig**. **(A)** Left panel: Pelleting test for F-actin (lanes 2, 3), α-actinin (lanes 5, 6) and BSA (lanes 8, 9), all in isolation at 21 °C. Right panel shows another example of F-actin pelleting test in isolation and the extent of actin found in the pellet fraction in this case (a faint band in the pellet lane). **(B)** F-actin pelleting control test by α-actinin and BSA at 21 °C. **(C)** F-actin pelleting control test by α-actinin and BSA at 4 °C. In both pelleting control experiments, the lanes are as follows: 1 (MW), 2 and 3 (F-actin + α-actinin), 5 and 6 (trial 1 of F-actin + BSA), 8 and 9 (trial 2 of F-actin + BSA). Centrifugation fraction labels on top of the gels are as follows: S, supernatant and P, pellet. The molecular weight (MW) marker values are in kDa.

**S9 Fig**. Pelleting of F-actin by Link-HP (done in triplicate) at 21 °C. Panel **(A)** shows trial 1 with Link-HP at 5.9 µM and **(B)** -trials 2, 3 of the same test. Panel **(C)** shows trial 1 with Link-HP at 1.6 µM and **(D)** -trials 2, 3 of the same test. Centrifugation fraction labels on top of the gels are as follows: S, supernatant and P, pellet. The molecular weight (MW) marker values are in kDa. The outcomes of all the controls are the same as for the VL147N tests at 5.9 µM (**Fig. 2** and **Fig. S8**).

**S10 Fig**. Pelleting of F-actin by Link-HP in presence of 100 mM KCl at 21 °C. Panel **(A)** shows trials 1 for Link-HP at 1.6 μM and 5.9 μM, **(B)** -trials 2, 3 for Link-HP at 5.9 μM and **(C)** -trials 2, 3 for Link-HP at 1.6 μM. Centrifugation fraction labels on top of the gels are as follows: S, supernatant and P, pellet. The molecular weight (MW) marker values are in kDa. The outcomes of all the controls are the same as for the VL147N tests at 5.9 µM (**Fig. 2** and **Fig. S8**).

**S11 Fig**. Pelleting of F-actin by VL147N in presence of 100 mM KCl at 21 °C. Panel **(A)** shows trials 1 for VL147N at 1.6 μM and 5.9 μM, **(B)** -trials 2, 3 for VL147N at 5.9 μM and **(C)** -trials 2, 3 for VL147N at 1.6 μM. Centrifugation fraction labels on top of the gels are as follows: S, supernatant and P, pellet. The molecular weight (MW) marker values are in kDa. The outcomes of all the controls are the same as for the VL147N tests at 5.9 µM (Fig. 2 and Suppl. Fig. 8).

**S12 Fig**. **(A)** Pelleting of F-actin by VHP101C (5.9 µM), trials 1 (lanes 3, 5) and 2 (lanes 7, 9), done at 4 °C. **(B)** Pelleting of F-actin by VHP101C (5.9 µM), trials 1 (lanes 3, 5) and 2 (lanes 7, 9), done in the presence of 100 mM KCl at 21 °C. When mixed with VHP101C, the bulk of F-actin remains in solution (supernatant) under all the temperature and [KCl] values tested. Centrifugation fraction labels on top of the gels are as follows: S, supernatant and P, pellet. The molecular weight (MW) marker values are in kDa. The outcomes of all the controls are the same as for the VL147N tests at 5.9 µM (**Fig. 2** and **Fig. S8**).

**S13 Fig**. Typical control experiments for F-actin bundling. **(A)** pelleting test of F-actin (lanes 3, 5) and BSA at 1.6 µM (lanes 7, 9) in isolation. **(B)** Pelleting of F-actin by BSA, trials 1 (lanes 3, 5) and 2 (lanes 7, 9). **(C)** Pelleting of F-actin by α-actinin. **(D)** Pelleting of α-actinin in isolation. All the tests shown here were done at 21 °C. Centrifugation fraction labels on top of the gels are as follows: S, supernatant and P, pellet. The molecular weight (MW) marker values are in kDa.

**S14 Fig**. VL147N at 3 °C without KCl (black contours) and in the presence of 150 mM KCl (red contours). Panels (**A**) and (**B**) show the same two spectra at different levels of minimal signal strength shown. Arrows show a pair of the related black and red signals which differ visibly in terms of their chemical shift values: Δ^1^H ≤ 0.03 ppm. Another pair of such peaks is shown in **Fig. 8**.

**S15 Fig**. VL147N at 25 °C without KCl (black contours) and in the presence of 150 mM KCl (red contours). Arrows show a pair of the related black and red signals which differ visibly in terms of their chemical shift values: Δ^1^H ≤ 0.03 ppm.

**S16 Fig**. Link-HP in the absence of KCl (black contours) and in the presence of 150 mM KCl (red contours) at 25 °C. Panel (**A**) shows the spectra in full, whereas the other panels show the same two spectra with the focus on different areas: (**B**) shows the bulk of the backbone ^1^H-^15^N resonances within the IDR parts of the samples and (**C**) focuses on the backbone ^1^H-^15^N signals of the glycine residues. Grey arrows indicate the signals (at least 8 instances) originating from Link-HP in the presence of 150 mM KCl which have no direct counterparts in the KCl-free spectrum.

**S17 Fig**. VL147N (black contours) and Link-HP (red contours) at 25 °C in the presence of 150 mM KCl. Panel (**A**) shows the bulk of the backbone ^1^H-^15^N resonances within the IDR parts of the samples and (**B**) focuses on the part of ^1^H-^15^N signals of the glycine residues where there are detected differences. Almost all the related black and red signals differ little in terms of their chemical shift values: Δ^1^H ≤ 0.02 ppm, Δ^15^N ≤ 0.07 ppm. Only in three instances (grey arrows), VL147N signals have no Link-HP counterparts, which can be explained by the covalent differences between the two samples at the C-terminus of VL147N. Also, in the vast majority of the cases, the relative intensities of the related VL147N (black contours) and Link-HP (red contours) signal differ little.

**S1 Table**. *Each (%Actin pelleted) value and standard deviation (std.dev.) represents the quantification of supernatant and pellet band intensities as estimated using ImageJ. All values, except for VHP101C, represent the average of three trials (n=3) ±standard deviation. The data for VHP101C were determined from 2 (n=2) measurements. The F-actin bundling outcomes form three non-overlapping groups: strong, intermediate and weak/absent (see **Fig. 5**). The strong bundling is characterized by high %F-actin pelleted with the minimal value equal 75% (as for 1.6 μM VL147N at 21 °C). The intermediate bundling outcomes has the maximal fraction of F-actin pelleted at 65% (as for 5.9 μM Link-HP at 21 °C) and minimal fraction at 32% (also for 5.9 μM Link-HP at 21 °C). The weak/absent bundling is characterized by the maximal fraction of F-actin pelleted of 9% (as for 5.9 μM VHP101C at 21 °C). The level of NaCl was 50 mM in all the samples tested.

## AUTHOR INFORMATION

### Author Contributions

Conceptualization: LJP, JMA, SLS

Data curation: SLS, LJP, JMA

Formal analysis: SKC, WPP, JMA, SLS

Funding acquisition: LJP, JMA, SLS

Investigation: SKC, WPP, JSH, LJP, JMA, SLS, CMH, EJC,

DLM Methodology: SKC, CMH, EJC, DLM, LJP, JMA, SLS

Project administration: SLS, JMA, LJP.

Resources: LJP, JMA, SLS

Supervision: JMA, SLS, LJP

Validation: SKC, WPP, JSH, LJP, JMA, SLS

Visualization: SKC, WPP, JSH, JMA, SLS

Writing – original draft: SKC, LJP, JMA, SLS

Writing – review & editing: All the authors.

### Notes

The authors have declared that no competing interests exist.

## ACKNOWLEDGMENT

We thank Dr. Thomas Unfried for their helpful suggestions on statistical analysis and multiple comparison corrections. We also thank Prof. Jeff Young for helpful discussions at various stages of this project. Caleb Howerton, Erich Walkenhauer, Melissa Oueld Es Cheikh, Kelly Vandamme and Raphael Marcenac contributed to protein samples expression and purification at the starting stages of the project. Finally, we would like to thank Sarina Kiesser (WWU) for help with recording mass spectrometry data.

## ABBREVIATIONS

VLN: villin
NMR: Nuclear Magnetic Resonance spectroscopy
F-actin: actin filaments
LC/QTOF: liquid chromatography quadrupole time of flight mass spectrometry

## PROTEIN SEQUENCES ACCESSION ID

Experimental protein samples described in the Results section of the manuscript (VL147N, Link-HP and VL101C, **Fig. 1**) are derived from *Arabidopsis thaliana* VILLIN4 (AtVLN4), isoform 2, UniProt accession O65570. Other plant villin variants referred to in this manuscript (**Suppl. Fig. 4**) and their UniProt accessions are as follows: AtVLN1 (F4ILN8), AtVLN2 (O81644), AtVLN3 (O81645) and AtVLN5 (Q9LVC6). Rabbit muscle actin used for F-actin bundling testing was commercially acquired from Cytoskeleton.com (cat. # AKL99).

## REFERENCES

1. Cao M, Zou X, Li C, Lin Z, Wang N, Zou Z, et al. An actin filament branching surveillance system regulates cell cycle progression, cytokinesis and primary ciliogenesis. Nat Commun. 2023;14(1):1687.

2. Pollard TD, Cooper JA. Actin, a central player in cell shape and movement. Science. 2009;326(5957):1208–12.

3. Pollard TD, Borisy GG. Cellular motility driven by assembly and disassembly of actin filaments. Cell. 2003;112(4):453–65.

4. Zhang R, Xu Y, Yi R, Shen J, Huang S. Actin cytoskeleton in the control of vesicle transport, cytoplasmic organization, and pollen tube tip growth. Plant physiology. 2023;193(1):9–25.

5. Li P, Day B. Battlefield Cytoskeleton: Turning the Tide on Plant Immunity. Mol Plant Microbe Interact. 2019;32(1):25–34.

6. Zhang W, Staiger CJ. Revising the Role of Cortical Cytoskeleton during Secretion: Actin and Myosin XI Function in Vesicle Tethering. Int J Mol Sci. 2021;23(1).

7. Khurana P, Henty JL, Huang S, Staiger AM, Blanchoin L, Staiger CJ. Arabidopsis VILLIN1 and VILLIN3 have overlapping and distinct activities in actin bundle formation and turnover. Plant Cell. 2010;22(8):2727–48.

8. Revenu C, Courtois M, Michelot A, Sykes C, Louvard D, Robine S. Villin severing activity enhances actin-based motility in vivo. Mol Biol Cell. 2007;18(3):827–38.

9. Friederich E, Vancompernolle K, Louvard D, Vandekerckhove J. Villin function in the organization of the actin cytoskeleton. Correlation of in vivo effects to its biochemical activities in vitro. J Biol Chem. 1999;274(38):26751–60.

10. Weber A, Northrop J, Bishop MF, Ferrone FA, Mooseker MS. Nucleation of actin polymerization by villin and elongation at subcritical monomer concentration. Biochemistry. 1987;26(9):2528–36.

11. Ge D, Pan T, Zhang P, Wang L, Zhang J, Zhang Z, et al. GhVLN4 is involved in multiple stress responses and required for resistance to Verticillium wilt. Plant science : an international journal of experimental plant biology. 2021;302:110629.

12. Wang X, Bi S, Wang L, Li H, Gao BA, Huang S, et al. GLABRA2 Regulates Actin Bundling Protein VILLIN1 in Root Hair Growth in Response to Osmotic Stress. Plant physiology. 2020;184(1):176–93.

13. Nag S, Larsson M, Robinson RC, Burtnick LD. Gelsolin: the tail of a molecular gymnast. Cytoskeleton (Hoboken). 2013;70(7):360–84.

14. Ferrary E, Cohen-Tannoudji M, Pehau-Arnaudet G, Lapillonne A, Athman R, Ruiz T, et al. In vivo, villin is required for Ca(2+)-dependent F-actin disruption in intestinal brush borders. J Cell Biol. 1999;146(4):819–30.

15. Trosch R. How villin subclasses coordinate actin remodelling. Nat Plants. 2024.

16. Li X, Zhuang Y, Zhao W, Qu X, Wang J, Chang M, et al. Molecular and functional adaption of Arabidopsis villins. The New phytologist. 2024.

17. Huang S, Qu X, Zhang R. Plant villins: versatile actin regulatory proteins. Journal of integrative plant biology. 2015;57(1):40–9.

18. Ghoshdastider U, Popp D, Burtnick LD, Robinson RC. The expanding superfamily of gelsolin homology domain proteins. Cytoskeleton (Hoboken). 2013;70(11):775–95.

19. Khurana S, George SP. Regulation of cell structure and function by actin-binding proteins: villin’s perspective. FEBS Lett. 2008;582(14):2128–39.

20. Miears HL, Gruber DR, Horvath NM, Antos JM, Young J, Sigurjonsson JP, et al. Plant Villin Headpiece Domain Demonstrates a Novel Surface Charge Pattern and High Affinity for F-Actin. Biochemistry. 2018;57(11):1690–701.

21. Boyko KV, Rosenkranz EA, Smith DM, Miears HL, Oueld Es Cheikh M, Lund MZ, et al. Sortase-mediated segmental labeling: A method for segmental assignment of intrinsically disordered regions in proteins. PLoS One. 2021;16(10):e0258531.

22. Zhuang Y, Wang Y, Jiao C, Shang Z, Huang S. Arabidopsis VILLIN5 bundles actin filaments using a novel mechanism. Plant J. 2024.

23. Altschul SF, Gish W, Miller W, Myers EW, Lipman DJ. Basic local alignment search tool. J Mol Biol. 1990;215(3):403–10.

24. Fedechkin SO, Brockerman J, Pfaff DA, Burns L, Webb T, Nelson A, et al. Gelsolin-like activation of villin: calcium sensitivity of the long helix in domain 6. Biochemistry. 2013;52(45):7890–900.

25. Chen L, Brown JW, Mok YF, Hatters DM, McKnight CJ. The allosteric mechanism induced by protein kinase A (PKA) phosphorylation of dematin (band 4.9). J Biol Chem. 2013;288(12):8313–20.

26. Rahmati Ishka M, Julkowska M. Tapping into the plasticity of plant architecture for increased stress resilience. F1000Res. 2023;12:1257.

27. Zhang H, Zhu J, Gong Z, Zhu JK. Abiotic stress responses in plants. Nat Rev Genet. 2022;23(2):104–19.

28. Wang C, Zhang LJ, Huang RD. Cytoskeleton and plant salt stress tolerance. Plant signaling & behavior. 2011;6(1):29–31.

29. Li M, Liu S, Wang J, Cheng X, Diao C, Yan D, et al. Dynamics of Actin Filaments Play an Important Role in Root Hair Growth under Low Potassium Stress in Arabidopsis thaliana. Int J Mol Sci. 2024;25(16).

30. Zhang P, Qian D, Luo C, Niu Y, Li T, Li C, et al. Arabidopsis ADF5 Acts as a Downstream Target Gene of CBFs in Response to Low-Temperature Stress. Front Cell Dev Biol. 2021;9:635533.

31. Lv F, Wang S, Tian R, Wang P, Liu K. Villin Family Members Associated with Multiple Stress Responses in Cotton. Phyton. 2021;90(6):1645–60.

32. Pacheco JM, Gabarain VB, Lopez LE, Lehuede TU, Ocaranza D, Estevez JM. Understanding signaling pathways governing the polar development of root hairs in low-temperature, nutrient-deficient environments. Current opinion in plant biology. 2023;75:102386.

33. Yuan G, Gao H, Yang T. Exploring the Role of the Plant Actin Cytoskeleton: From Signaling to Cellular Functions. Int J Mol Sci. 2023;24(20).

34. Kumar S, Jeevaraj T, Yunus MH, Chakraborty S, Chakraborty N. The plant cytoskeleton takes center stage in abiotic stress responses and resilience. Plant, cell & environment. 2023;46(1):5–22.

35. Zamora-Briseno JA, Pereira-Santana A, Reyes-Hernandez SJ, Cerqueda-Garcia D, Castano E, Rodriguez-Zapata LC. Towards an understanding of the role of intrinsic protein disorder on plant adaptation to environmental challenges. Cell Stress Chaperones. 2021;26(1):141–50.

36. Abdul Aziz M, Sabeem M, Mullath SK, Brini F, Masmoudi K. Plant Group II LEA Proteins: Intrinsically Disordered Structure for Multiple Functions in Response to Environmental Stresses. Biomolecules. 2021;11(11).

37. Choura M, Rebai A, Hanin M. Proteome-wide analysis of protein disorder in Triticum aestivum and Hordeum vulgare. Comput Biol Chem. 2020;84:107138.

38. Covarrubias AA, Cuevas-Velazquez CL, Romero-Perez PS, Rendon-Luna DF, Chater CCC. Structural disorder in plant proteins: where plasticity meets sessility. Cell Mol Life Sci. 2017;74(17):3119–47.

39. Holehouse AS, Das RK, Ahad JN, Richardson MO, Pappu RV. CIDER: Resources to Analyze Sequence-Ensemble Relationships of Intrinsically Disordered Proteins. Biophys J. 2017;112(1):16–21.

40. Das RK, Pappu RV. Conformations of intrinsically disordered proteins are influenced by linear sequence distributions of oppositely charged residues. Proc Natl Acad Sci U S A. 2013;110(33):13392–7.

41. Smirnov SL, Isern NG, Jiang ZG, Hoyt DW, McKnight CJ. The Isolated Sixth Gelsolin Repeat and Headpiece Domain of Villin Bundle F-Actin in the Presence of Calcium and Are Linked by a 40-Residue Unstructured Sequence. Biochemistry. 2007;46(25):7488–96.

42. Wilkins MR, Gasteiger E, Bairoch A, Sanchez JC, Williams KL, Appel RD, et al. Protein identification and analysis tools in the ExPASy server. Methods Mol Biol. 1999;112:531–52.

43. 43. Walker JM. The Proteomics Protocols Handbook. 2005:571–607.

44. Anthis NJ, Clore GM. Sequence-specific determination of protein and peptide concentrations by absorbance at 205 nm. Protein Sci. 2013;22(6):851–8.

45. Csizmok V, Follis AV, Kriwacki RW, Forman-Kay JD. Dynamic Protein Interaction Networks and New Structural Paradigms in Signaling. Chem Rev. 2016;116(11):6424–62.

46. DeForte S, Uversky VN. Order, Disorder, and Everything in Between. Molecules. 2016;21(8).

47. Dyson HJ. Making Sense of Intrinsically Disordered Proteins. Biophys J. 2016;110(5):1013–6.

48. Bah A, Forman-Kay JD. Modulation of Intrinsically Disordered Protein Function by Post-translational Modifications. J Biol Chem. 2016;291(13):6696–705.

49. Martin EW, Holehouse AS, Grace CR, Hughes A, Pappu RV, Mittag T. Sequence Determinants of the Conformational Properties of an Intrinsically Disordered Protein Prior to and upon Multisite Phosphorylation. J Am Chem Soc. 2016;138(47):15323–35.

50. Das RK, Ruff KM, Pappu RV. Relating sequence encoded information to form and function of intrinsically disordered proteins. Curr Opin Struct Biol. 2015;32:102–12.

51. Trosch R. How villin subclasses coordinate actin remodelling. Nat Plants. 2025;11(1):6.

52. Fedechkin SO, Brockerman J, Luna EJ, Lobanov MY, Galzitskaya OV, Smirnov SL. An N-terminal, 830-residue Intrinsically Disordered Region of the Cytoskeleton-regulatory Protein Supervillin Contains Myosin II-and F-actin-Binding Sites. J Biomol Struct Dyn. 2013;31(10):1150–9.

53. Crowley JL, Smith TC, Fang Z, Takizawa N, Luna EJ. Supervillin reorganizes the actin cytoskeleton and increases invadopodial efficiency. Mol Biol Cell. 2009;20(3):948–62.

54. Chen Y, Takizawa N, Crowley JL, Oh SW, Gatto CL, Kambara T, et al. F-actin and myosin II binding domains in supervillin. J Biol Chem. 2003;278(46):46094–106.

55. Blotnick E, Sol A, Muhlrad A. Histones bundle F-actin filaments and affect actin structure. PLoS One. 2017;12(8):e0183760.

56. Huang C, Ni Y, Wang T, Gao Y, Haudenschild CC, Zhan X. Down-regulation of the filamentous actin cross-linking activity of cortactin by Src-mediated tyrosine phosphorylation. J Biol Chem. 1997;272(21):13911–5.

57. Gifford JL, Jamshidiha M, Mo J, Ishida H, Vogel HJ. Comparing the calcium binding abilities of two soybean calmodulins: towards understanding the divergent nature of plant calmodulins. Plant Cell. 2013;25(11):4512–24.

58. Thomas C. Bundling actin filaments from membranes: some novel players. Frontiers in plant science. 2012;3:188.

59. Yang H, Wei X, Wang L, Zheng P, Li J, Zou Y, et al. Functional Characterization of PeVLN4 Involved in Regulating Pollen Tube Growth from Passion Fruit. Int J Mol Sci. 2025;26(5).

60. Qu X, Zhang H, Xie Y, Wang J, Chen N, Huang S. Arabidopsis villins promote actin turnover at pollen tube tips and facilitate the construction of actin collars. Plant Cell. 2013;25(5):1803–17.

61. Zhang Y, Xiao Y, Du F, Cao L, Dong H, Ren H. Arabidopsis VILLIN4 is involved in root hair growth through regulating actin organization in a Ca2+-dependent manner. The New phytologist. 2011;190(3):667–82.

62. Zou M, Ren H, Li J. An Auxin Transport Inhibitor Targets Villin-Mediated Actin Dynamics to Regulate Polar Auxin Transport. Plant physiology. 2019;181(1):161–78.

